# c-di-GMP inhibits early sporulation in *Clostridioides difficile*

**DOI:** 10.1101/2021.06.24.449855

**Authors:** Adrianne N. Edwards, Caitlin L. Willams, Nivedita Pareek, Shonna M. McBride, Rita Tamayo

**Affiliations:** Department of Microbiology and Immunology, Emory University School of Medicine, Emory Antibiotic Resistance Center, Atlanta, Georgia, United States of America; Department of Microbiology and Immunology, University of North Carolina Chapel Hill, Chapel Hill, North Carolina, United States of America

**Keywords:** *Clostridioides difficile*, *Clostridium difficile*, sporulation, spore, cyclic diguanylate, c-di-GMP, anaerobe, cyclic diguanylate synthase

## Abstract

The formation of dormant spores is essential for the anaerobic pathogen *Clostridioides difficile* to survive outside of the host gastrointestinal tract. The regulatory pathways and environmental signals that initiate *C. difficile* spore formation within the host are not well understood. One bacterial second messenger signaling molecule, cyclic diguanylate (c-di-GMP), modulates several physiological processes important for *C. difficile* pathogenesis and colonization, but the impact of c-di-GMP on sporulation is unknown. In this study, we investigated the contribution of c-di-GMP to *C. difficile* sporulation. Overexpression of a gene encoding a diguanylate cyclase, *dccA*, decreased sporulation frequency and early sporulation gene transcription in both the epidemic R20291 and historical 630Δ*erm* strains. Expression of a *dccA* allele encoding a catalytically inactive DccA that is unable to synthesize c-di-GMP no longer inhibited sporulation, indicating that the accumulation of intracellular c-di-GMP reduces *C. difficile* sporulation. A null mutation in *dccA* slightly increased sporulation in R20291 and slightly decreased sporulation in 630Δ*erm*, suggesting that DccA may contribute to the intracellular pool of c-di-GMP in a strain-dependent manner. However, these data were highly variable, underscoring the complex regulation involved in modulating intracellular c-di-GMP concentrations. Finally, overexpression of *dccA* in known sporulation mutants revealed that c-di-GMP is likely signaling through an unidentified regulatory pathway to control early sporulation events in *C. difficile*. C-di-GMP-dependent regulation of *C. difficile* sporulation may represent an unexplored avenue of potential environmental and intracellular signaling that contributes to the complex regulation of sporulation initiation.

**IMPORTANCE:** Many bacterial organisms utilize the small signaling molecule cyclic diguanylate (c-di-GMP) to regulate important physiological processes, including motility, toxin production, biofilm formation, and colonization. C-di-GMP inhibits motility and toxin production and promotes biofilm formation and colonization in the anaerobic, gastrointestinal pathogen *Clostridioides difficile*. However, the impact of c-di-GMP on *C. difficile* spore formation, a critical step in this pathogen’s life cycle, is unknown. Here, we demonstrate that c-di-GMP negatively impacts sporulation in two clinically relevant *C. difficile* strains, the epidemic R20291 and the historical 630Δ*erm*. The pathway through which c-di-GMP controls sporulation was investigated, and our results suggest that c-di-GMP is likely signaling through an unidentified regulatory pathway to control *C. difficile* sporulation. This work implicates c-di-GMP metabolism as a potential mechanism to integrate environmental and intracellular cues through c-di-GMP levels to influence *C. difficile* sporulation.

## INTRODUCTION

Nucleotide second messengers, such as the nearly ubiquitous cyclic diguanylate (c-di-GMP), serve as central intracellular signaling molecules in bacteria. C-di-GMP promotes the switch from a planktonic, motile stage to a sessile, surface-associated lifestyle and controls virulence factor production in numerous pathogenic and nonpathogenic bacteria. C-di-GMP is synthesized from GTP by diguanylate cyclases (DGCs, synthases), which contain the conserved catalytic GGDEF motif (1). Phosphodiesterases (PDEs, hydrolases) containing either the EAL or HD-GYP domain degrade c-di-GMP to pGpG or GMP, respectively (2–5). Often, DGC and PDE proteins contain additional sensory or regulatory domains that potentially control enzymatic activity, suggesting environmental and bacterial cues influence the intracellular concentration of c-di-GMP (6). Most Gram-negative bacteria encode numerous DGC and PDE proteins, resulting in complex c-di-GMP metabolic pathways, while Gram-positive bacteria often contain a more modest number of DGC and PDE proteins (7–9). However, the gastrointestinal pathogen *Clostridioides difficile* encodes many c-di-GMP metabolic proteins; 37 genes encoding GGDEF and/or EAL domains were identified in the historical 630 strain (10, 11). Many of the *C. difficile* c-di-GMP metabolic proteins have been demonstrated to be enzymatically active (10, 12), suggesting that the regulation of c-di-GMP metabolism in *C. difficile* is physiologically important and tightly controlled. High intracellular concentrations of c-di-GMP have been demonstrated to inhibit *C. difficile* motility and toxin production while promoting cell aggregation, biofilm formation, and colonization (13–20).

*C. difficile* is an obligate anaerobe and relies on the formation of a dormant spore for long-term persistence outside of the host and transmission to new hosts. Spore formation in all endospore-forming bacteria, including *C. difficile*, is initiated by the highly conserved transcriptional regulator Spo0A (21, 22). Spo0A activity is tightly controlled by phosphorylation through a large regulatory network of kinases, phosphatases, and additional regulators (23, 24). Once phosphorylated, Spo0A~P activates the expression of early sporulation genes, triggering sporulation (25). However, many of the regulatory proteins and pathways that control early sporulation events in the model organism *Bacillus subtilis* are not conserved in *C. difficile* (26, 27). Although recent progress has uncovered several important sporulation regulatory factors in *C. difficile* (reviewed in 28), the environmental cues and regulatory pathways that control Spo0A activation are largely unknown.

Environmental conditions and nutrient availability likely trigger *C. difficile* spore formation within the host gastrointestinal tract. Two global transcriptional regulators, CodY and CcpA, control sporulation initiation in *C. difficile* in response to nutrient availability. CodY represses target gene expression when GTP and branched-chain amino acids are abundant, and CcpA activates or represses target gene transcription based on carbohydrate availability (29, 30). Mutations in either CodY or CcpA result in increased sporulation frequencies (30, 31). Some sporulation genes serve as direct targets for CodY and CcpA regulation, but the molecular mechanisms are not delineated (29–31). Additionally, the uptake of peptides, a critical nutrient source for *C. difficile* (32),by the Opp and App oligopeptide permeases affects the timing of sporulation, as *opp* and *app* inactivation significantly increases sporulation frequency (33). It is reasonable to consider that other global signaling systems in *C. difficile* link nutrient availability and/or other environmental conditions to the decision to initiate sporulation.

In this study, we examined the impact that c-di-GMP has on *C. difficile* sporulation. We showed that overexpression of *dccA*, a gene encoding a DGC, resulted in decreased sporulation in two important *C. difficile* strains. The conserved catalytic motif, GGDEF, was required for DccA-dependent inhibition of *C. difficile* spore formation, indicating that the c-di-GMP metabolic activity is responsible for this phenotype. Finally, we provided evidence that c-di-GMP does not depend on signaling through several known sporulation factors, suggesting that c-di-GMP influences sporulation through an unidentified pathway.

## RESULTS

### Overexpression of a diguanylate cyclase reduces *C. difficile* sporulation frequency

As c-di-GMP affects many physiological processes in *C. difficile*, we hypothesized that *C. difficile* sporulation is also influenced by c-di-GMP. To test this hypothesis, *dccA*, which encodes a DGC in *C. difficile* (10, 13), was overexpressed on a multicopy plasmid using the nisin-inducible *cpr* promoter (13, 33, 34) in two different *C. difficile* backgrounds. We included R20291, which is a clinically-prevalent epidemic strain, and 630Δ*erm*, which is a spontaneous erythromycin-sensitive derivative of 630, a clinical isolate which has served as a long-term laboratory model strain (35, 36). Of note, the amino acid sequences of DccA from R20291 and 630Δ*erm* are 100% identical. To assess sporulation frequency, we performed ethanol-resistant sporulation assays in these strains after 24 h of growth on 70:30 sporulation agar.

When a plasmid copy of *dccA* (pDccA) was overexpressed from the nisin-inducible promoter in the R20291 background, sporulation frequency significantly decreased from 33.7% in the absence of nisin to 19% in the presence of 0.5 μg/ml nisin (**Fig. 1A**), suggesting that overexpression of *dccA*, and presumably high intracellular levels of c-di-GMP, inhibit *C. difficile* sporulation. To assess whether the decrease in sporulation frequency was due to the production of c-di-GMP by DccA, we overexpressed a *dccA* allele encoding a mutated GGDEF domain (AADEF) that is unable to synthesize c-di-GMP (pDccAmut; 13). Here, even at the highest expression of *dccA* (0.5 μg/ml nisin), sporulation efficiency remained unaffected, indicating that the DccA-dependent reduction in sporulation frequency is due to DccA’s diguanylate cyclase activity. We also visualized sporulation in these strains using phase contrast microscopy in which spores appear phase-bright and vegetative cells are phase-dark rods. When *dccA* was overexpressed in R20291, not only were fewer spores visible, but this strain also formed long chains (**Fig. 1B**). This cell morphology change was also dependent on increased intracellular concentrations of c-di-GMP, as this effect was not seen in the R20291 strain overexpressing pDccA^mut^ (data not shown). This change in cell morphology with overexpression of *dccA* was expected and is likely due to c-di-GMP-dependent increase in expression of *cmrRST*, which encodes an atypical signal transduction system that regulates *C. difficile* colony morphology and motility (37).

**Figure 1.**
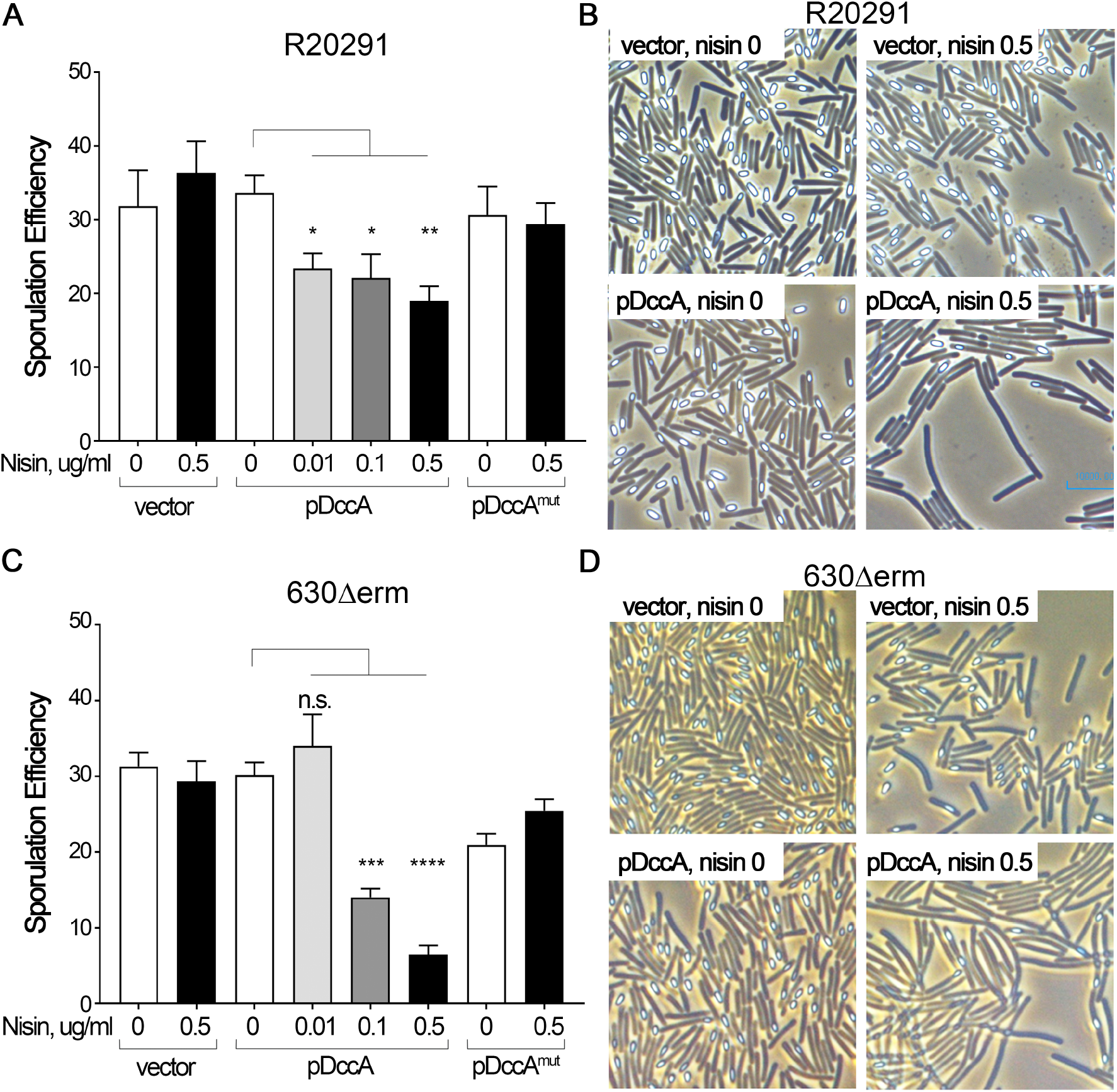
Overexpression of *dccA* inhibits sporulation in R20291 and 630Δ*erm* and is dependent upon a functional cyclic diguanylate (GGDEF) domain. Ethanol-resistant sporulation assays (**A**) and representative phase contrast micrographs (**B**) of R20291 pMC211 (RT526), R20291 pDccA (RT527), and R20291 pDccA^mut^ (RT539) grown on 70:30 sporulation agar supplemented with 2 μg/ml thiamphenicol and 0-0.5 μg/m nisin as indicated at H_24_. Ethanol-resistant sporulation assays (**C**) and representative phase contrast micrographs (**D**) of 630Δ*erm* pMC211 (RT762), 630Δ*erm* pDccA (RT763), and 630Δ*erm* pDccA^mut^ (RT764) grown on 70:30 sporulation agar supplemented with 2 μg/ml thiamphenicol and 0-0.5 μg/m nisin as indicated at H_24_. The means and standard errors of the means for at least three biological replicates are shown. *p<0.05, **p<0.01, ***p<0.001, ****p<0.0001, one-way ANOVA with Dunnett’s post-test comparing values to uninduced WT pDccA.

When *dccA* was overexpressed in the 630Δ*erm* background, we observed an ~4-fold decrease in sporulation efficiency (29.7% in the absence of nisin to 8.5% in 0.5 μg/ml nisin; **Fig. 1C**). This reduction in sporulation frequency in the 630Δ*erm* pDccA strain was dose-dependent, with the lowest sporulation frequency occurring in the highest expression of *dccA* (0.5 μg/ml nisin). Again, overexpression of the *dccA*^mut^ allele resulted in wild-type levels of sporulation, indicating that the ability of DccA to repress sporulation relies on a functional GGDEF domain and its diguanylate cyclase activity. Similar to R20291, overexpression of *dccA* in the 630Δ*erm* background resulted in fewer spores and a change in cell morphology when observed by phase contrast microscopy, and these phenotypes were also dependent on a functional DccA GGDEF domain (**Fig. 1D**; data not shown).

### Overexpression of a cyclic diguanylate decreases sporulation-specific gene expression

To ensure that *dccA* and *dccA*^mut^ transcription are activated in a dose-dependent manner in increasing concentrations of nisin, we measured *dccA* transcript levels using qRT-PCR. Cells were harvested after 12 h of growth on 70:30 sporulation agar, which marks early stationary phase and the onset of sporulation (38). As expected, *dccA* and *dccA*^mut^ transcript levels were increased ~4-6-fold in both R20291 and 630Δ*erm* in the presence of 0.5 μg/ml nisin (**Fig. 2A, B**), showing that *dccA* and *dccA*^mut^ expression are consistent in both backgrounds. To better understand the effect of c-di-GMP on early sporulation events in *C. difficile*, we measured the steady-state transcript levels of an early sporulation-specific gene, *sigE* (*spoIIG*), using qRT-PCR. Transcription of *sigE* is dependent on active, phosphorylated Spo0A (39). The relative expression of *sigE* in both R20291 pDccA and 630Δ*erm* pDccA grown on 0.5 μg/ml nisin was decreased ~2-fold compared to their respective parent strains (**Fig. 2C, D**). Decreased *sigE* transcript levels in R20291 pDccA were dependent upon a functional GGDEF domain as *sigE* transcript levels were unchanged when *dccA^mut^* was overexpressed in R20291 (**Fig. 2C**). These data indicate that overexpression of *dccA* reduces Spo0A-dependent gene expression and confirm that the increased production of c-di-GMP is responsible for reduced sporulation in *C. difficile*.

**Figure 2.**
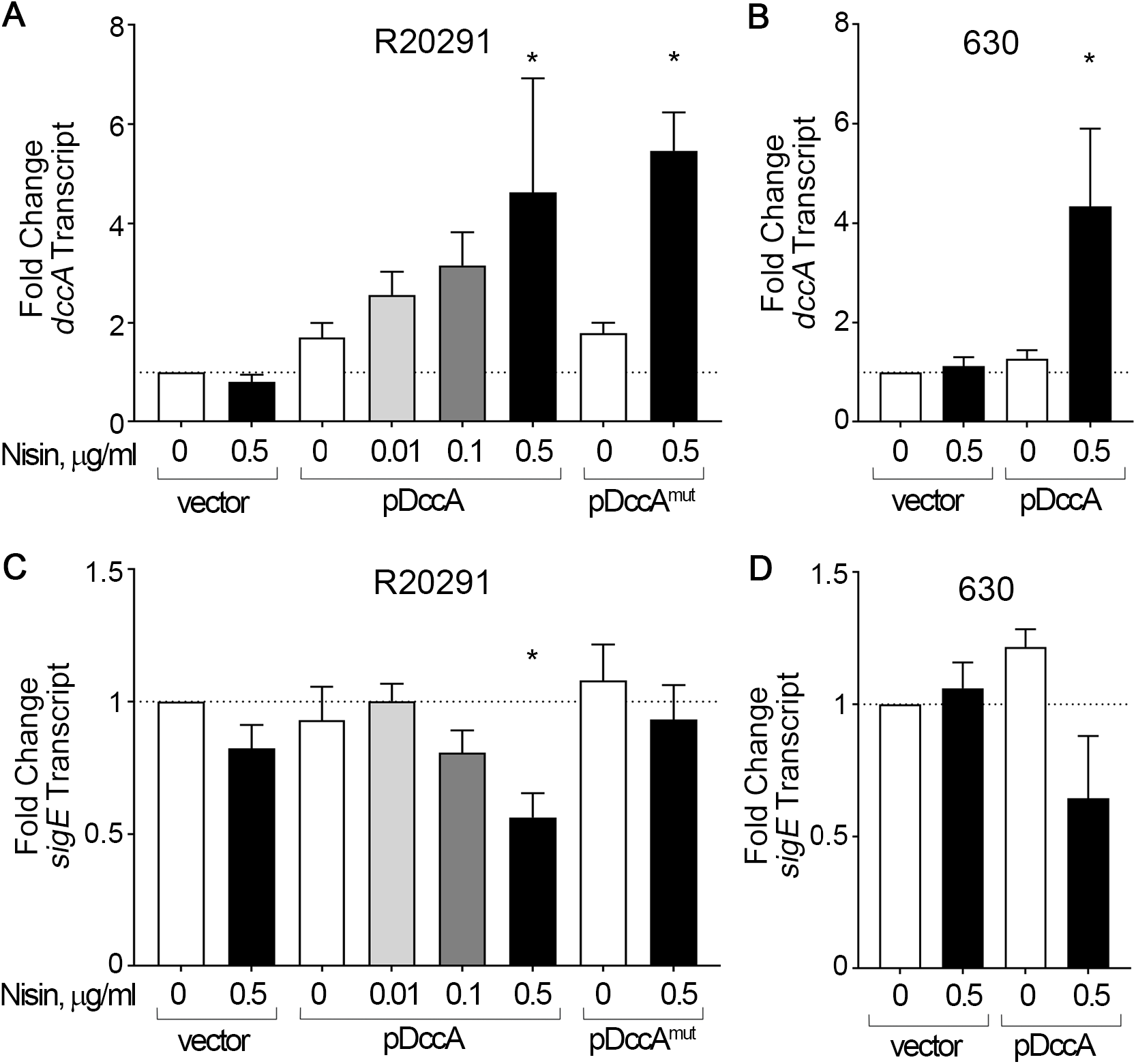
Overexpression of *dccA* decreases Spo0A-dependent gene expression. qRT-PCR analyses of *dccA* (**A**) and *sigE* (**C**) transcript levels in R20291 pMC211 (RT526), R20291 pDccA (RT527), and R20291 pDccA^mut^ (RT539) grown on 70:30 sporulation agar supplemented with 2 μg/ml thiamphenicol and 0-0.5 μg/m nisin as indicated at H_12_. qRT-PCR analyses of *dccA* (**B**) and *sigE* (**D**) transcript levels in 630Δ*erm* pMC211 (RT762) and 630Δ*erm* pDccA (RT763) grown on 70:30 sporulation agar supplemented with 2 μg/ml thiamphenicol and in the absence or presence of 0.5 μg/ml nisin at H_12_. The means and standard errors of the means for at least three biological replicates are shown. *p<0.05, **p<0.01, one-way ANOVA with Dunnett’s post-test comparing values to uninduced WT vector. *p<0.05, **p<0.01.

### Deletion of *dccA* results in variable sporulation frequencies in R20291 and 630Δ*erm*

Sporulation may be impacted by one or a subset of c-di-GMP metabolic enzymes. *C. difficile* 630 encodes 37 proteins containing GGDEF and/or EAL domains, and R20291 encodes 31 (10, 11, 35). Overexpression of *dccA* bypasses endogenous control of c-di-GMP. To better understand the c-di-GMP regulatory mechanism, we next asked whether a null mutation in *dccA* alone affects *C. difficile* sporulation frequency. To test this hypothesis, we employed the pseudo-suicide vectors, pMSR and pMSR0 (tailored for the 630 and R20291 backgrounds respectively), which take advantage of allelic exchange and an inducible *C. difficile* toxin-antitoxin system to create a markerless gene deletion (40).

The R20291 Δ*dccA* mutant produced slightly more spores than R20291 (42.3% in the R20291 Δ*dccA* versus 31.7% in the isogenic parent; **Fig. 3A**). However, the sporulation frequencies were highly variable, and the differences between strains were not statistically significant. The R20291 Δ*dccA* mutant was complemented by integrating the *dccA* gene into the chromosome using the conjugative transposon, Tn916. *dccA* is the second gene in a two gene operon in R20291 and 630Δ*erm*, and the promoter of the upstream gene, *CD1421*, was used to drive *dccA* transcription in the complementation construct. The sporulation frequency of R20291 Δ*dccA* Tn*916*∷P_*CD1421*_-*dccA* was reduced (34.7%) compared to the *dccA* mutant, but again, the data were variable and not statistically significant.

**Figure 3.**
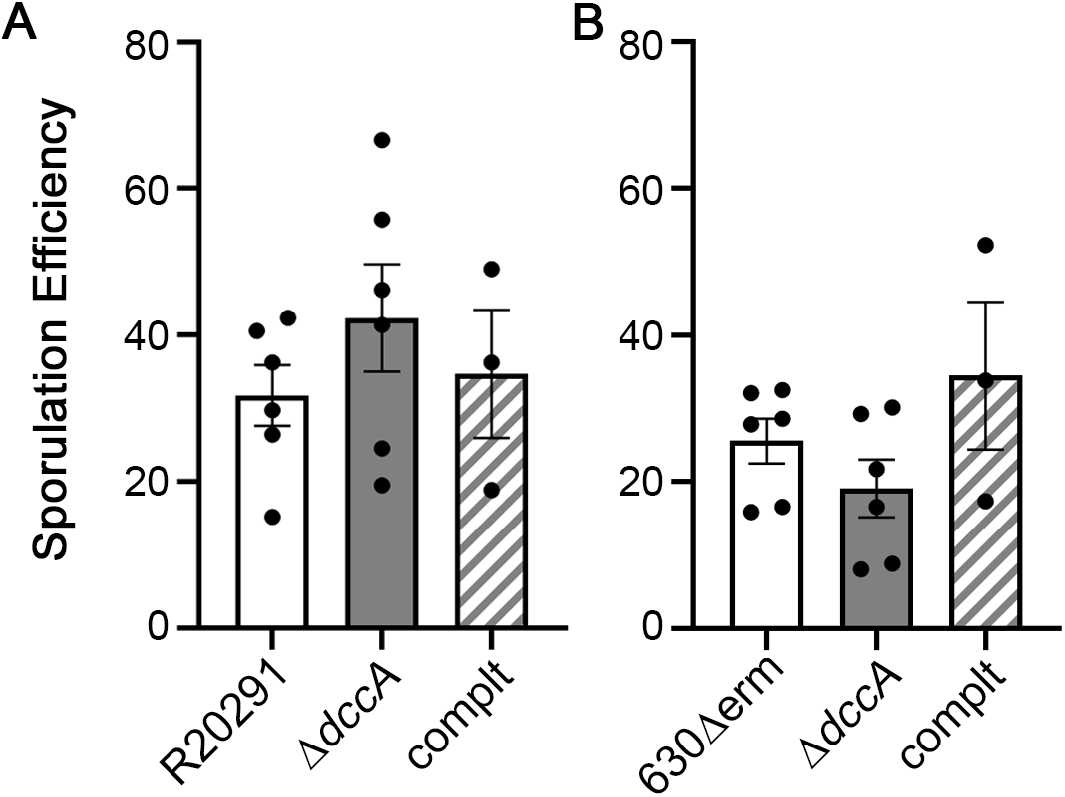
A null *dccA* mutation variably affects R20291 and 630Δ *erm* sporulation. Ethanol-resistant sporulation assays of (**A**) R20291, R20291 Δ*dccA* (RT2656), and R20291 Δ*dccA* Tn*916*∷P_*CD1421*_-*dccA* (MC1961) and (**B**) 630Δ*erm*, 630Δ*erm* Δ*dccA* (RT2703), and 630Δ*erm* Δ*dccA* Tn*916*∷P_*CD1421*_-*dccA* (MC1960) grown on 70:30 sporulation agar at H_24_. The means and standard errors of the means for at least three biological replicates are shown. No significant differences by one-way ANOVA.

The 630Δ*erm* Δ*dccA* mutant exhibited a slightly reduced sporulation frequency compared to the 630Δ*erm* parent (19.1% in the *dccA* mutant versus 25.6% in the parent strain; **Fig. 3B**), an opposite trend compared to the R20291 Δ*dccA* mutant. The complemented strain, 630Δ*erm* Δ*dccA* Tn*916*∷P_*CD1421*_-*dccA*, showed an increased sporulation frequency (34.4%). But, as in the R20291 background, the sporulation frequencies were variable and did not achieve statistical significance. Altogether, these data suggest that c-di-GMP synthesis by DccA does not greatly and/or consistently contribute to sporulation initiation in the conditions tested, or that other c-di-GMP metabolic enzymes are redundant with or compensate for the loss of DccA.

### C-di-GMP likely does not signal through known *C. difficile* sporulation factors

To attempt to identify the regulatory pathway(s) through which c-di-GMP influences sporulation in *C. difficile*, we overexpressed *dccA* in several well-studied 630Δ*erm* sporulation mutants and performed ethanol-resistant sporulation assays after 24 h growth on 70:30 sporulation agar. We included the parent 630Δ*erm* strain overexpressing *dccA* in the absence and presence of 0.5 μg/ml nisin as a reference in these experiments (**Fig. 4A**).

**Figure 4.**
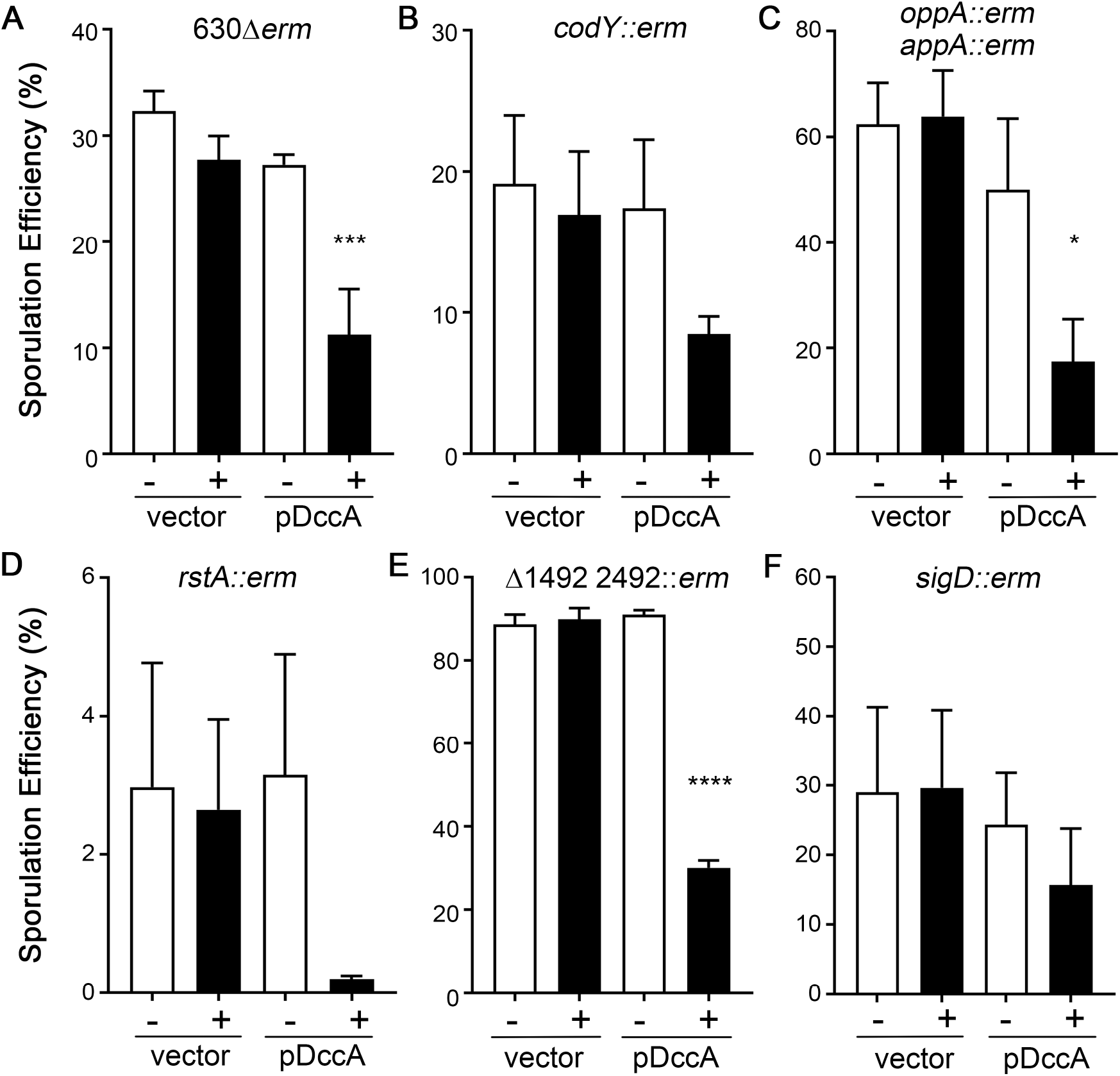
C-di-GMP does not inhibit sporulation through known sporulation factors. Ethanol-resistant sporulation assays of (**A**) 630Δ*erm* pMC211 (RT762) and 630Δ*erm* pDccA (RT763), (**B**) 630Δ*erm codY*∷*erm* pMC211 (MC947) and 630Δ*erm codY*∷*erm* pDccA (MC948), (**C**) 630Δ*erm oppB*∷*erm appA*∷*erm* pMC211 (MC924) and 630Δ*erm oppB*∷*erm appA*∷*erm* pDccA (MC925), (**D**) 630Δ*erm rstA*∷*erm* pMC211 (MC926) and 630Δ*erm rstA*∷*erm* pDccA (MC927), (**E**) 630Δ*erm* Δ*CD1492 CD2492*∷*erm* pMC211 (MC928) and 630Δ*erm* Δ*CD1492 CD2492*∷*erm* pDccA (MC929), and (**F**) 630Δ*erm sigD*∷*erm* pMC211 (MC864) and 630Δ*erm sigD*∷*erm* pDccA (MC865) grown on 70:30 sporulation agar supplemented with 2 μg/ml thiamphenicol and in the absence or presence of 0.5 μg/ml nisin as indicated at H_24_. The means and standard errors of the means for at least three biological replicates are shown. *p<0.05, ***p<0.001, ****p<0.0001, one-way ANOVA with Dunnett’s post-test comparing values to respective parent strain with vector, uninduced. Note that the y-axes for each panel vary depending on the sporulation frequency of the strain tested.

First, we assessed the effect of *dccA* overexpression in 630Δ*erm codY* and 630Δ*erm opp app* mutants. CodY is a global transcriptional regulator that binds to target DNA in high intracellular concentrations of the effectors GTP and branched-chain amino acids (BCAAs; 41). Loss of GTP and BCAA binding changes the conformation of CodY, differential expression of metabolic pathways and other physiological processes, including toxin production and sporulation (29, 31, 42, 43). The *opp* and *app* operons each encode oligopeptide permeases which import small, heterogenous peptides into the cell (44, 45). Inactivation of these permeases in *C. difficile* results in significantly increased sporulation, suggesting that limited nutrient uptake triggers sporulation (33). The regulatory pathways and molecular mechanisms by which CodY, Opp, and App affect sporulation are not fully understood, although null mutations in these loci affect sporulation timing and result in increased sporulation frequencies (31, 33). The sporulation frequencies of the *codY* mutant and the *opp app* double mutant decreased when *dccA* was overexpressed using 0.5 μg/ml nisin (2-fold and 3.7-fold, respectively, compared to each strain’s vector controls grown in 0.5 μg/ml nisin; **Fig. 4B, C**). These results indicate that c-di-GMP does not require CodY or the Opp and App oligopeptide permeases to inhibit *C. difficile* sporulation.

Next, we ascertained whether RstA or the phosphotransfer proteins, CD1492 and CD2492, are part of the regulatory pathway c-di-GMP employs to control sporulation. RstA is a multifunctional protein that serves as a transcriptional regulator and, through a separate domain, positively influences *C. difficile* sporulation initiation via an unknown mechanism (38). Overexpression of *dccA* in the *rstA* background resulted in an ~13-fold decreased sporulation frequency compared to the *rstA* mutant containing the vector control (**Fig. 4D**), indicating that RstA is not necessary for c-di-GMP-dependent inhibition of sporulation. CD1492 and CD2492 are predicted histidine kinases that function as phosphotransfer proteins to repress sporulation and are hypothesized to directly impact Spo0A phosphorylation (28, 46). A *CD1492 CD2492* double mutant exhibited significantly increased sporulation compared to the 630Δ*erm* parent (**Fig. 4A, E**), and overexpression of *dccA* in the *CD1492 CD2492* background reduced sporulation frequency ~3-fold (**Fig. 4E**), demonstrating that CD1492 and CD2492 are not a required part of the c-di-GMP signaling pathway.

Finally, we asked whether c-di-GMP signals through SigD to control *C. difficile* sporulation. SigD is the flagellar alternative sigma factor that is required for both motility and efficient toxin production in *C. difficile* (14, 38, 47). Although a null *sigD* mutation does not affect sporulation in these conditions (comparing 630Δ*erm* and 630Δ*erm sigD* vector control strains, **Fig. 4A, F** and (38, 47)), we chose to investigate the *sigD* mutant because c-di-GMP directly represses *sigD* transcription, inhibiting *C. difficile* motility and toxin production (13, 15, 48). Sporulation frequency was reduced only 1.9-fold when *dccA* was overexpressed in the *sigD* background, and the difference was not statistically significant (**Fig. 4F**). The DccA-dependent effect is not as dramatic as in the other mutant backgrounds tested; however, these data suggest that c-di-GMP does not primarily influence sporulation through SigD. Altogether, these data indicate that c-di-GMP does not significantly impact *C. difficile* sporulation through these known regulatory factors in the conditions tested.

## DISCUSSION

In this work, we set out to determine the impact of the bacterial second messenger c-di-GMP on *C. difficile* sporulation. We found that overexpression of *dccA*, encoding a DGC that synthesizes high intracellular levels of c-di-GMP when overexpressed (13), resulted in a significant decrease in early sporulation gene expression and spore formation in the epidemic R20291 and historical 630Δ*erm* strains. The conserved catalytic GGDEF motif was required for DccA-dependent inhibition of sporulation, indicating that the diguanylate cyclase synthase activity of DccA is responsible for decreased sporulation. Consistent with this result, sporulation was inhibited in a dose-dependent manner, as higher transcript levels of *dccA* coincided with fewer detected transcripts of *sigE*, which encodes an early sporulation-specific sigma factor.

Because DccA overexpression drastically altered the sporulation frequency in R20291 and 630Δ*erm*, we had anticipated a stronger sporulation phenotype in the corresponding *dccA* mutants. However, there are many additional DGCs encoded in the *C. difficile* genome, as well as many PDEs, and these likely contribute to the intracellular concentration of c-di-GMP also, as most are catalytically active (10). Given the sheer number of encoded DGCs and that their functions may inherently exhibit redundancy, it is, in retrospect, unsurprising that the deletion of a single GGDEF domain protein does not significantly impact sporulation. Any contribution DccA exerts on the intracellular c-di-GMP pool may be masked by the redundant functions of similar proteins. As such, we previously found that overexpression of DccA resulted in modest increases in transcript levels of several encoded PDEs, suggesting that *C. difficile* is able to somewhat compensate for high levels of intracellular c-di-GMP (49). It is also possible that changes in c-di-GMP levels upon loss of DccA are compensated for by the upregulation or activation of other DGCs, downregulation or inhibition of PDEs, or both. Importantly, previous studies have shown that distinct DGCs control different c-di-GMP-regulated phenotypes, suggesting that localized pools of c-di-GMP influence only a subset of phenotypes (50, 51). It is plausible that the deletion of one or more of the other encoded DGCs in *C. difficile* could result in a stronger impact on sporulation frequency. During preparation of this manuscript, a bioRxiv submission reported that overexpression of a phosphodiesterase containing an EAL domain increases sporulation, and deletion of that PDE decreases sporulation in *C. difficile* UK1, an epidemic strain that is nearly identical to R20291, corroborating our findings (52).

Interestingly, two early *C. difficile* sporulation regulators that are orthologous to the *B. subtilis* SinR transcriptional repressor regulate *C. difficile* intracellular c-di-GMP levels. Null mutations in the two *C. difficile* SinR orthologs, known as SinRR’ in R20291 and CD2214-CD2215 in 630Δ*erm*, result in increased *dccA* transcription and c-di-GMP levels, and an asporogenous phenotype (20, 53). Further, deletion of *CD2214-CD2215* resulted in differential expression of additional DGC and PDE genes in *C. difficile* 630Δ*erm* (20). It will be intriguing to determine if the effect SinRR’ on *C. difficile* sporulation occurs through alteration of c-di-GMP levels by regulating DGC/PDE gene expression or if SinRR’ affects sporulation initiation through multiple regulatory pathways.

Identification of the c-di-GMP effector(s) that mediate c-di-GMP-dependent sporulation regulation in *C. difficile* is of great interest. A variety of c-di-GMP receptors that directly bind to c-di-GMP have been characterized in bacteria. These encompass a number of protein-based receptors, including proteins containing degenerate GGDEF and/or EAL domains, and two distinct types of riboswitches that alter downstream gene expression in response to c-di-GMP binding (8). *C. difficile* encodes a single PilZ domain protein (6), a domain that often directly binds c-di-GMP (54, 55), and a Type IV pilus PilB ATPase similar to orthologs that have been shown to bind c-di-GMP (56), but there are no other known or predicted c-di-GMP protein receptors reported in *C. difficile* (8)*. C. difficile* encodes at least eleven functional riboswitches that alter gene expression in response to c-d-GMP and contains five additional predicted riboswitches (15, 49). None of these riboswitches appear to affect expression of sporulation-related genes; however, the conditions used in this study do not support efficient sporulation initiation in *C. difficile* (49). The regulatory pathway c-di-GMP utilizes to influence sporulation remains unknown. The decrease in spore formation mediated by *dccA* overexpression does not appear to signal through CodY, the Opp or App oligopeptide permeases, RstA, or the CD1492 and CD2492 phosphotransfer proteins. Although the effect of DccA-mediated inhibition of sporulation was slightly decreased in the *sigD* mutant, c-di-GMP unlikely signals solely through SigD in these conditions. Given that we know c-di-GMP directly affects *sigD* transcription in *C. difficile* through the Cdi-1-3 riboswitch (13, 15, 48), it is attractive to hypothesize that SigD might have a role in sporulation. However, there has been no published evidence of a regulatory role between SigD and sporulation thus far (38, 47, 49).

The impact of c-di-GMP on sporulation has been investigated in only a few other spore-forming bacteria. Interestingly, using a mCherry reporter fused to a c-di-GMP-regulated riboswitch, high c-di-GMP levels were observed in sporulating *B. subtilis* cells, suggesting a correlation (57), but the impact of c-di-GMP in *B. subtilis* sporulation has remained relatively unexplored. Studies in *Bacillus thuringiensis* and *Bacillus anthracis* found no effect on sporulation efficiency when any of the catalytically active GGDEF/EAL/HD-GYP encoding genes were individually deleted (58, 59). These studies in *Bacillus* sp. further underscore the differences between *C. difficile* and other endospore-forming bacteria in their early sporulation regulatory networks. Finally, direct c-di-GMP regulation of sporulation has only been identified in *Streptomyces* sp., where c-di-GMP inhibits spore formation directly by antagonizing the sporulation-specific sigma factor, WhiG, and binding directly to the transcriptional regulator, BldD (60, 61). The contribution of c-di-GMP to sporulation remains an understudied field.

Utilizing c-di-GMP to inhibit sporulation may be advantageous to *C. difficile*, as c-di-GMP can be rapidly degraded when environmental and intracellular conditions favor sporulation. The c-di-GMP metabolic activity of a protein is often controlled by the corresponding domains encoded within the same protein. Identifying the DGCs and PDEs that affect *C. difficile* sporulation and investigating the function of their associated domains may reveal the environmental and intracellular signals that promote or delay sporulation. Finally, the finding that c-di-GMP is a regulator of early sporulation events in *C. difficile* creates new research opportunities for discovering the potentially novel regulatory pathways, c-di-GMP effectors, and molecular mechanisms that control spore formation in this significant pathogen.

## MATERIALS AND METHODS

### Bacterial strains and growth conditions

The bacterial strains and plasmids used for this study are listed in **Table 1**. *Clostridioides difficile* was routinely cultured in BHIS medium in a 37°C vinyl anaerobic chamber (Coy) with an atmosphere of 5% CO_2_, 10% H_2_, and 85% N_2_ (62). *C. difficile* cultures were supplemented with 2-10 μg/ml thiamphenicol as necessary for plasmid maintenance. *Escherichia coli* strains were grown aerobically at 37°C in LB with 100 μg/ml ampicillin and/or 10-20 μg/ml chloramphenicol, and counterselection against *E. coli* after conjugation with *C. difficile* was performed using 100 μg/ml kanamycin (13).

**Table 1.**
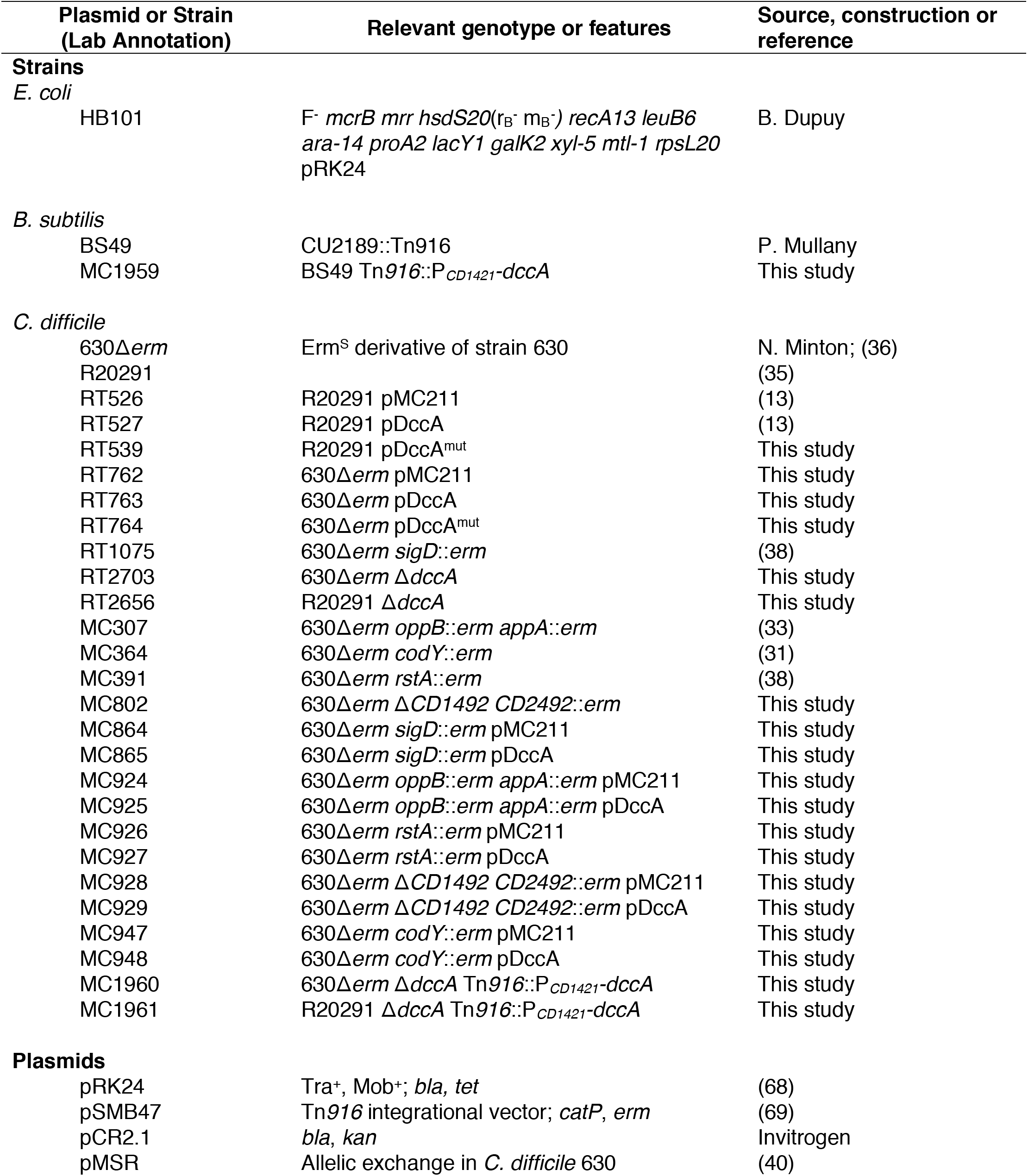
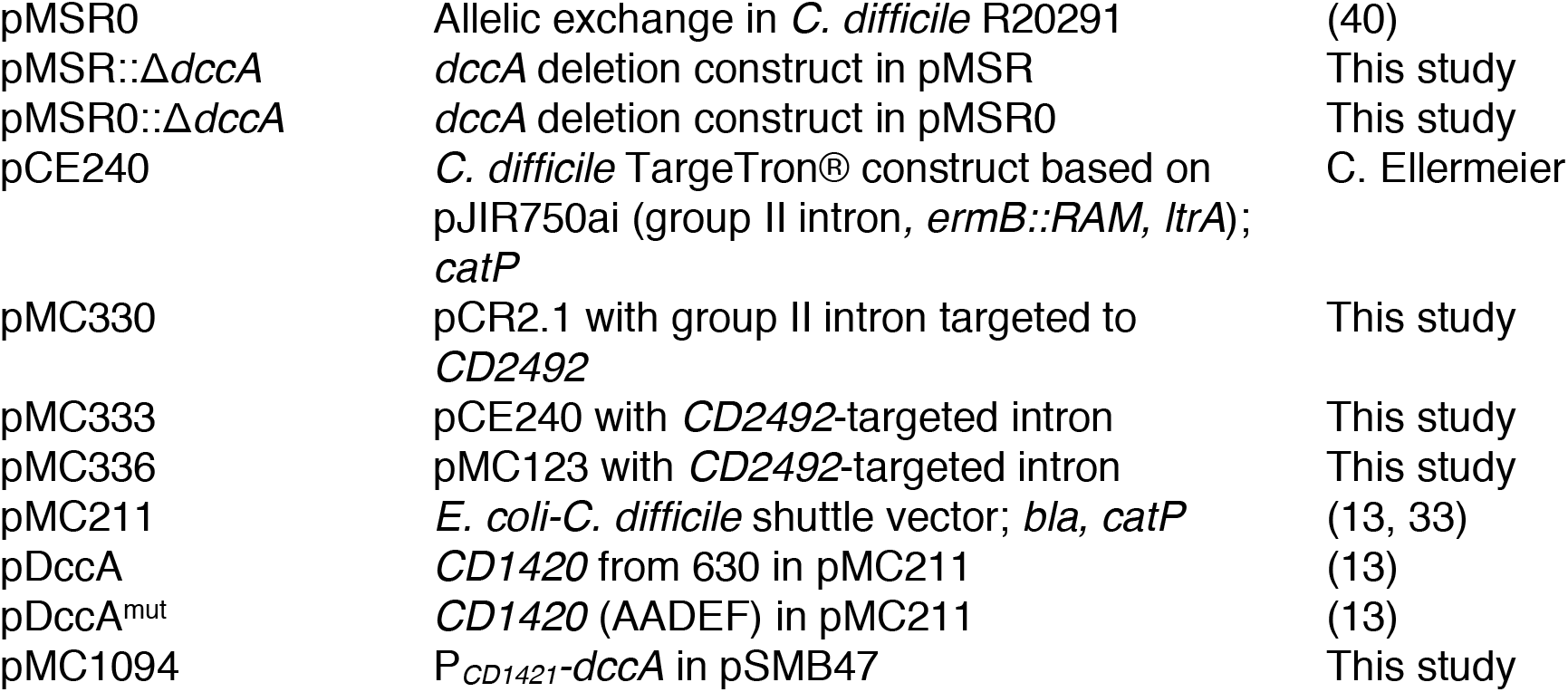
Bacterial Strains and plasmids.

**Table 2.**
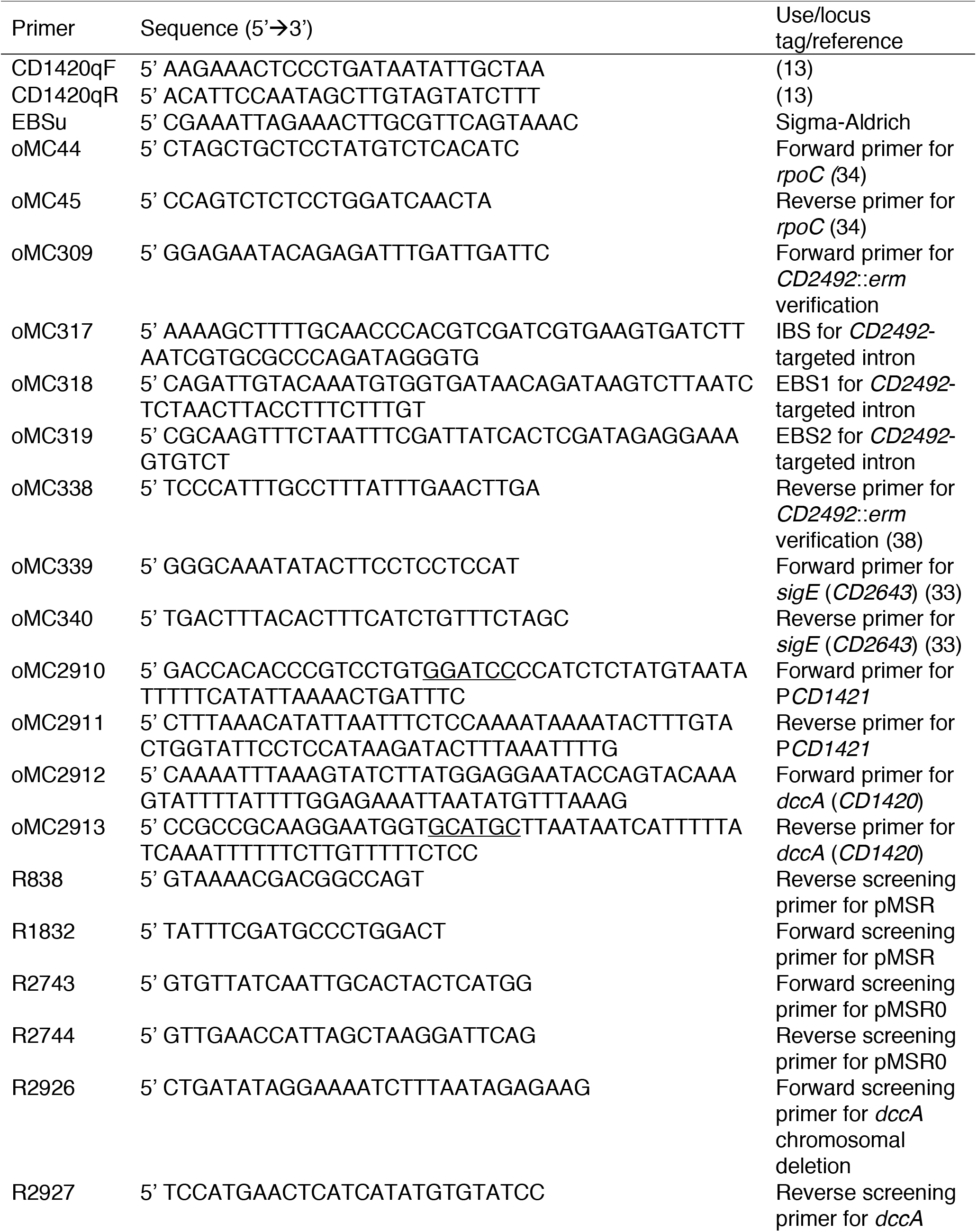
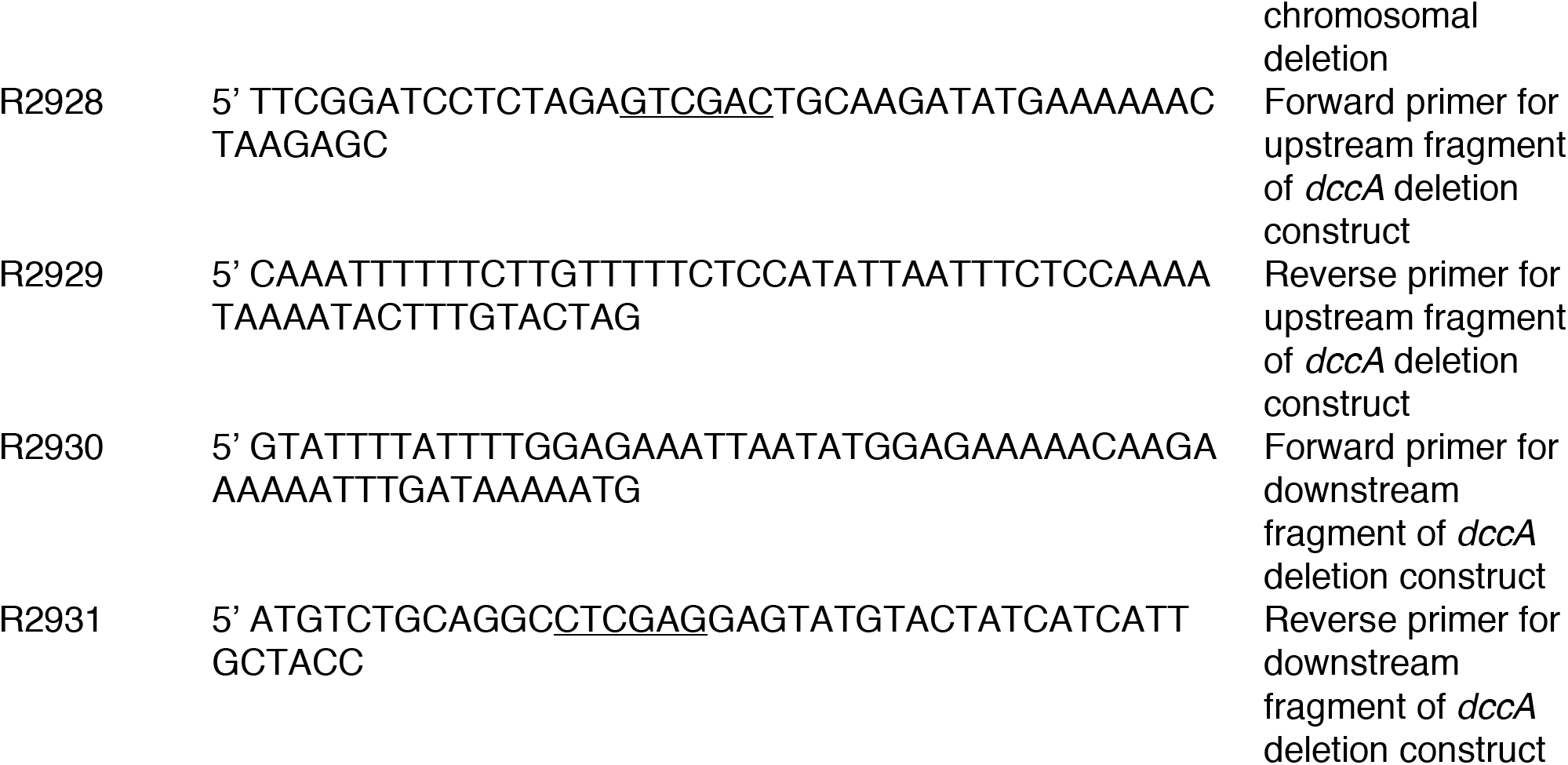
Oligonucleotides.

### Strain and plasmid construction

*C. difficile* strains 630 (GenBank accession no. NC_009089.1) and R20291 (GenBank accession no. FN545816.1) were used as the basis for primer design and PCR amplification. The *dccA* mutants were constructed using the pseudo-suicide vectors pMSR and pMSR0 (40). Upstream and downstream homology regions were amplified from the 630Δ*erm* genome with primers R2928/R2929 and R2930/R2931, respectively. The fragments were Gibson assembled (NEB) into *Sal*I/*Xho*I-digested pMSR to create pMSR∷Δ*dccA*. Similar fragments were amplified from the R20291 chromosome using the same primers and assembled into pMSR0 to create pMSR0∷Δ*dccA*. Chloramphenicol-resistant colonies were confirmed by PCR with plasmid-specific primers R838/R1832 (pMSR) or R2743/R2744 (pMSR0).

To create the *dccA* mutants, the pMSR∷Δ*dccA* and pMSR0∷Δ*dccA* plasmids were transformed into *E. coli* HB101 pRK24 for conjugation with *C. difficile* 630Δ*erm* and R20291, respectively. Subsequent steps were done essentially as previously described (40). Briefly, large thiamphenicol- and kanamycin-resistant colonies, which presumably had integrated the plasmid into the chromosome to allow for more rapid growth, were streaked again on BHIS with 10 μg/ml thiamphenicol and 100 μg/ml kanamycin to ensure purity. Then large colonies were streaked on BHIS with 100 ng/ml anhydrotetracycline (ATc) to induce expression of the toxin gene and eliminate bacteria that still contained the vector. Colonies were screened for the 0.8 kb deletion of *dccA* using primers R2926/R2927 (**Fig. 5A**). Genomic DNA was isolated from potential mutants, and the *dccA* region was PCR amplified using primers R2926/R2927 and sequenced to confirm integrity of the sequence.

**Figure 5.**
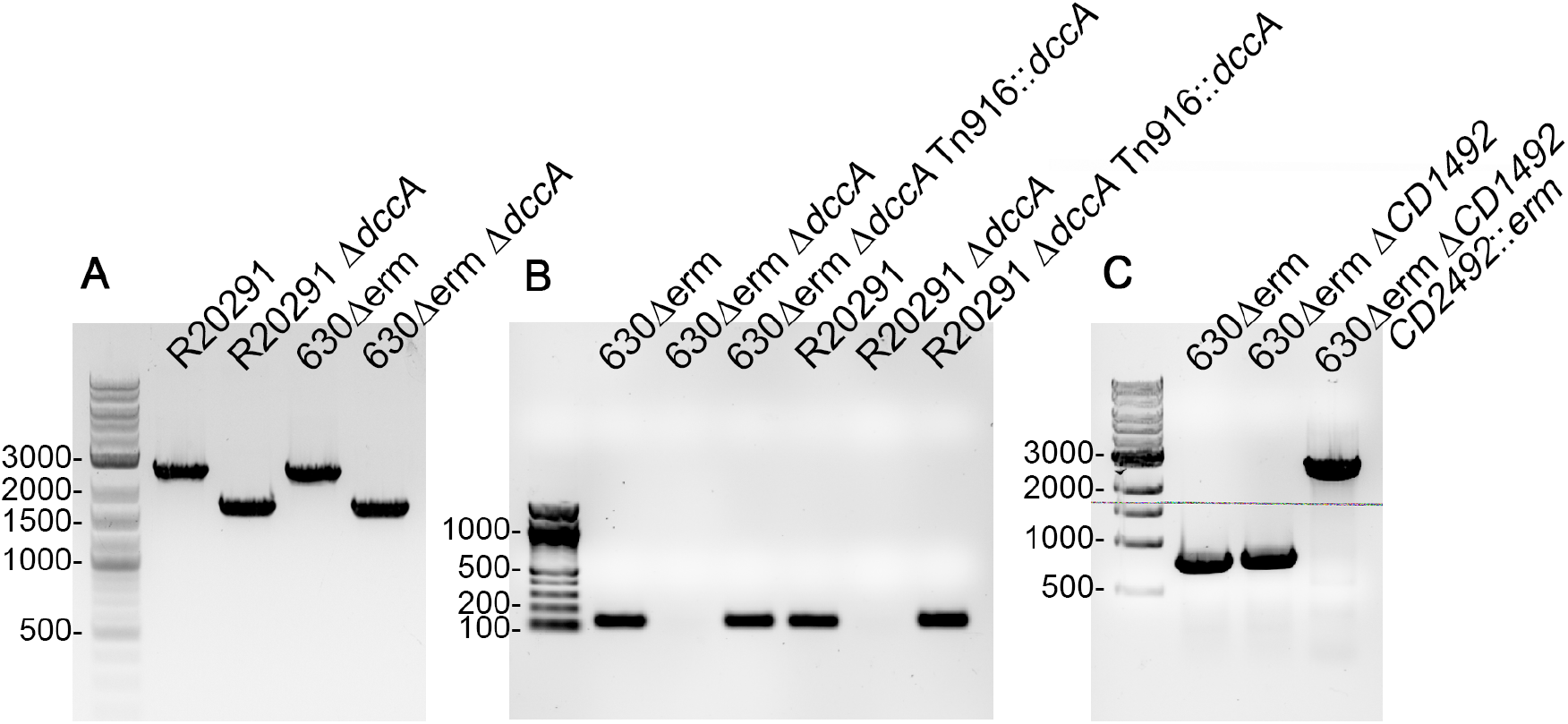
PCR verification of constructed strains. (**A**) PCR confirmation of Δ*dccA* in R20291 and 630Δ*erm* backgrounds using primers R2926/R2927. Shown are strains R20291, R20291 Δ*dccA* (RT2656), 630Δ*erm*, and 630Δ*erm* Δ*dccA* (RT2703). Expected PCR products’ sizes are 2.5 kb for the wild-type allele and 1.7 kb for the deletion mutants. (**B**) PCR verification of Tn*916*∷P_*CD1421*_-*dccA* chromosomal integration in 630Δ*erm* and R20291 using the internal qRT-PCR *dccA* primers CD1420qF/CD1420qR. Shown are strains 630Δ*erm* Δ*dccA* (RT2703), 630Δ*erm* Δ*dccA* Tn*916*∷P_*CD1421*_-*dccA* (MC1960), R20291, R20291 Δ*dccA* (RT2656), and R20291 Δ*dccA* Tn*916*∷P_*CD1421*_-*dccA* (MC1961). Expected PCR product size is 140 bp. (**C**) PCR confirmation of *CD2492*-targeted intron in the 630Δ*erm* Δ*CD1492* background (MC674). Shown are strains 630Δ*erm*, 630Δ*erm* Δ*CD1492* (MC674), and 630Δ*erm* Δ*CD1492 CD2492*∷*erm* (MC802) using primers oMC309/oMC338. The expected PCR products’ sizes are 811 bp for the wild-type *CD2492* allele and ~2811 bp for the *CD2492*∷*erm*.

The 630Δ*erm* and R20291 *dccA* mutants were complemented using a *Bacillus subtilis* donor strain, BS49, carrying the conjugative transposon, Tn916, to transfer in the *dccA* gene driven by its native promoter, which is encoded upstream of *CD1421* (MC1959). To create the plasmid carrying the Tn*916*∷P_*CD1421*_-dccA construct (pMC1094), P_*CD1421*_ and *dccA* were spliced by overlapping PCR using primers oMC2910/2911 and oMC2912/2913 and Gibson assembled into *Bam*HI/*Sph*I-digested pSMB47 as. Erythromycin-resistant colonies were confirmed by PCR with primers CD1420qF/CD1420qR (**Fig. 5B**).

To create the 630Δ*erm* Δ*CD1492 CD2492*∷*erm* double mutant, the Targetron-based group II intron from pCE240 was retargeted using the previously published targeting site by Underwood *et. al*. 2009 to create pMC336 (63). Briefly, the *CD2492*-targeted intron was amplified using primers oMC317, oMC318, oMC319 and EBSu and TA-cloned into pCR2.1 to create pMC330. A *BsrG*I/*Hind*III-digested fragment containing the *CD2492-* targeted intron was subcloned into pCE240 to create pMC333. Finally, a *Sph*I/*Sfo*I-digested fragment containing the *CD2492-*targeted intron was subcloned into pMC123 to create pMC336. The resulting pMC336 was conjugated into the 630Δ*erm CD1492* strain (MC674) and erythromycin-resistant mutants were screened for the 2 kb Targetron insertion within *CD2492* using primers oMC309/338 (**Fig. 5C**). Notably, the targeting site was not located in the 254a site within the CD2492 coding region noted in Underwood *et al*. 2009 (63), but rather at 318s (data not shown).

### Sporulation assays

*C. difficile* strains were grown overnight in BHIS medium supplemented with 0.1% taurocholate and 0.2% fructose to aid in spore germination and to prevent spore formation, respectively (64, 65). Thiamphenicol (5 μg/ml) was included for plasmid maintenance when necessary. When strains reached mid-exponential phase (OD_600_ ≅ 0.5), 250 μl aliquots were applied to the surface of 70:30 agar containing 2 μg/ml thiamphenicol and 0-0.5 μg/ml nisin (65). After 24 hours of growth, ethanol-resistant sporulation assays were performed as previously described (46, 66). Briefly, cells were scraped from the plate surface and suspended in BHIS to an OD_600_ ≅ 1. To enumerate vegetative cells, the cell suspensions were serially diluted in BHIS and plated onto BHIS plates. Simultaneously, 0.5 ml aliquots of the cell suspensions were mixed thoroughly with 0.3 ml 95% ethanol and 0.2 ml dH_2_O (final concentration = 28.5% ethanol) and incubated for 15 min to eliminate all vegetative cells. Ethanol-treated cells were then serially diluted in 1X PBS with 0.1% taurocholate and plated onto BHIS supplemented with 0.1% taurocholate. Total colony forming units (CFU) were enumerated after at least 24 h of growth, and the sporulation frequency was calculated as the number of ethanol-resistant spores divided by the total number of vegetative and ethanol-resistant spores combined. A *spo0A* mutant (MC310) was used as a negative control in all assays.

### Phase contrast microscopy

Phase contrast microscopy was performed at H_24_ as described previously (33). Briefly, cells were concentrated by pelleting 0.5 ml of the cell suspension, and the concentrated cell suspension was applied to a 0.7% agarose pad on a slide. Cells were imaged with a 100X Ph3 oil immersion objective on a Nikon Eclipse Ci-L microscope. At least two fields of view were captured with a DS-Fi2 camera from at least three independent experiments for each strain tested.

### Quantitative reverse transcription PCR analysis (qRT-PCR)

RNA was isolated from *difficile* strains grown on 70:30 sporulation agar at H_12_ and DNase-I treated as previously described (13, 29, 33, 34). cDNA was synthesized using random hexamers, and quantitative real-time-PCR (qRT-PCR) reactions were performed in triplicate (Bioline) and monitored by a Roche Lightcycler 96. The *rpoC* transcript (oMC44/oMC45) was used as the reference gene (34). Controls with no reverse transcriptase were included for all templates and primer sets. The data were analyzed by the 2^−ΔΔCT^ method (67), with normalization to *rpoC* and the stated reference condition or strain. The results represent the means and standard error of the means of at least three independent experiments.

## ACKNOWLEDGEMENTS

We are grateful to the members of the Tamayo and McBride labs for helpful suggestions and discussions during the course of this work. This research was supported by the U.S. National Institutes of Health through research grants AI107029 and AI143638 to R.T. and AI116933 and AI156052 to S.M.M. C.L.W. is supported by an Institutional Research and Academic Career Development Awards (IRACDA) fellowship under K12-GM000678. The content of this manuscript is solely the responsibility of the authors and does not necessarily reflect the official views of the National Institutes of Health.

